# Common Signatures of Neutrophils in Diverse Disease Conditions

**DOI:** 10.1101/2025.07.02.662891

**Authors:** Di Wu, Ying Cao, Hongjie Chen, Minghao Du, Tingting Wang, Jiarui Zhao, Mengyuan Li, Wenjing Wu, Huixuan Zhang, Ence Yang, Jing Yang, Jian Chen

## Abstract

Neutrophils are traditionally recognized for their pro-inflammatory roles. However, accumulating evidence has begun to highlight the plasticity of transcriptional states of neutrophils during pathological insults. Whether such unconventional neutrophils may share commonality across diverse disease conditions is incompletely understood. Here, we systematically profile over 500,000 neutrophils in key immune compartments and disease-inflicted tissues of the mouse models of metabolic disorder, autoimmunity, tissue damage, peripheral cancers, and intracranial gliomas. Of importance, we observe two distinct neutrophil clusters with unique immune features. The first cluster, characterized by Cd274, Vegfa, and antigen presentation, is highly enriched within the diseased tissues. In contrast, the second cluster with elevated Cd244 and type 2 immune response but reduced maturation markers primarily emerges in the peripheral blood and spleens of specific disease scenarios. These results have elucidated the common signatures of neutrophils in response to various pathological conditions, providing a valuable reference for diagnostic and therapeutic applications.

## Main Text

Neutrophils are the most abundant immune cells in the body and can account for over 50% of white blood cells. It has long been documented that neutrophils respond to various pathological insults, e.g., infection or tissue damage, by phagocytosis, releasing pro-inflammatory factors (i.e., degranulation), or forming neutrophil extracellular traps (i.e., NETosis) ^1,2^. Notably, in contrast to those conventional roles, recent studies based on single-cell RNA sequencing (scRNA-seq) by colleagues and us have begun to define various transcriptional states of neutrophils in different cancers, e.g., liver cancer, lung cancer, breast cancer, pancreatic cancer, and glioma ^3–8^. However, whether such unconventional neutrophils may commonly emerge under different disease conditions is incompletely understood.

We sought to systematically profile neutrophils in mouse disease models, i.e., metabolic disorder (high-fat diet, HFD), autoimmunity (experimental autoimmune encephalomyelitis, EAE), tissue damage (acute lung injury, ALI), peripheral cancers [Lewis lung carcinoma (LLC), B16 melanoma, and MC38 colorectal carcinoma], and intracranial tumors (LCPNS or LCPNS-SIIN gliomas ^8^). Neutrophils were FACS-sorted from the key immune compartments, i.e., bone marrow, spleen, and peripheral blood, and diseased tissues, i.e., liver, spinal cord, lung, or tumors, of those mouse models and pooled for scRNA-seq profiling (Table S1). We note that neutrophils from individual mice were not tracked in those scRNA-seq data. After the quality control, a total of 534,689 neutrophils were obtained and integrated into a scRNA-seq dataset (Fig. 1a), though potential batch effects might still remain a limitation. Among the defined transcriptional states, the marker of progenitor cells *Cd34*, the immature neutrophil marker *Camp*, and the proliferation markers *Mki67* decreased as granulocyte-monocyte progenitors (GMP) differentiated into more mature populations (Fig. 1b and Fig. S1a). Meanwhile, an overall increasing trend of classic neutrophil markers *Cxcr2* and *S100a8* occurred with neutrophil maturation (Fig. 1b). This putative differentiation path could be visualized by the RNA velocity analysis (Fig. 1c) and the pseudotime trajectory analysis (Fig. 1d), and those neutrophil transcriptional states had the distinct enrichment of the targeted genes of specific transcription factors (Fig. S1b). Notably, the distribution of neutrophil transcriptional states exhibited a profound difference across the diseased tissues (Fig. 1e and Fig. S2a) and the immune compartments, i.e., peripheral blood (Fig. 1f), spleens (Fig. S2b), and bone marrow (Fig. S2c).

**Figure 1.**
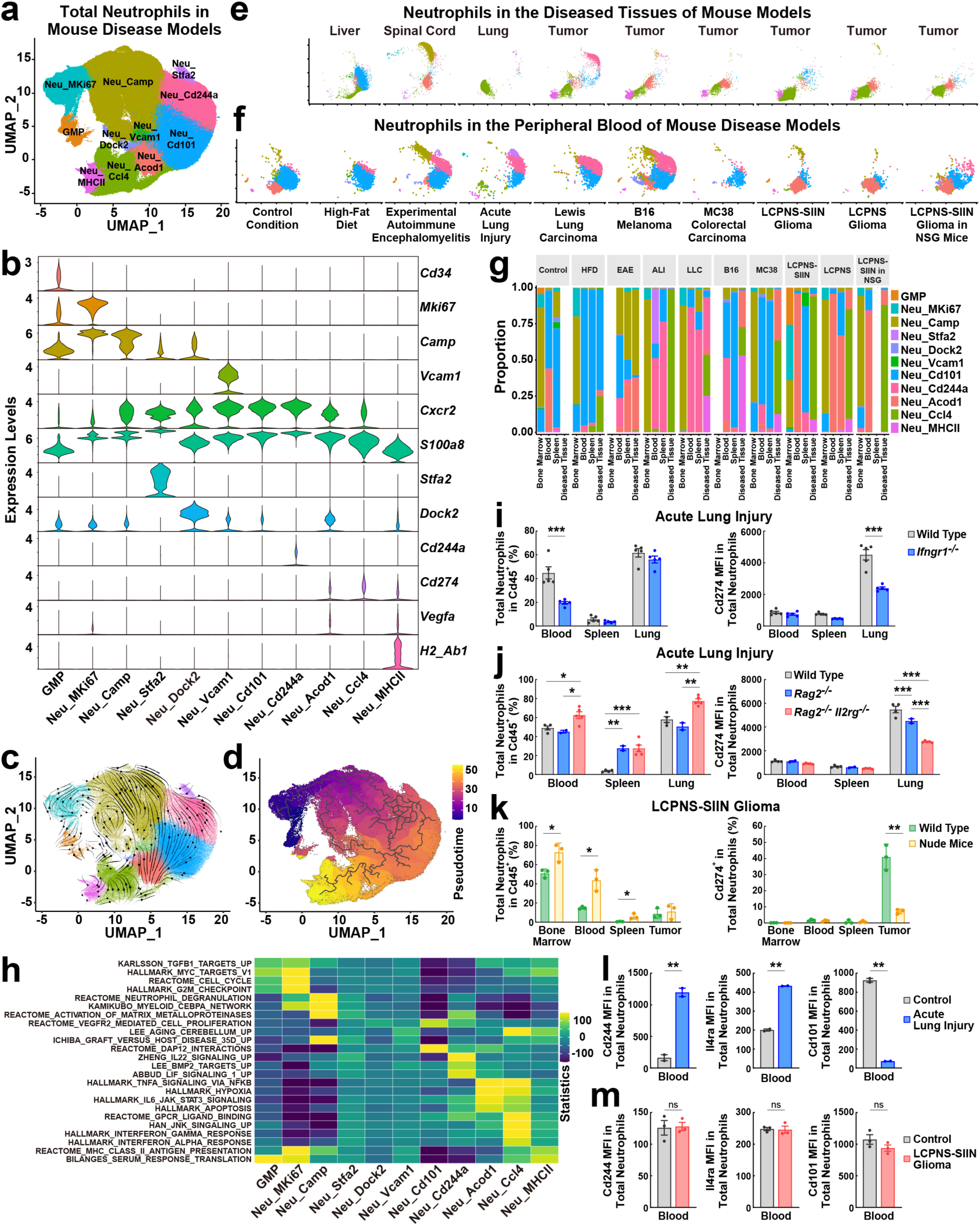
Common signatures of neutrophils across different disease conditions. **a** UMAP plot of neutrophil transcriptional states in the pooled scRNA-seq dataset of the bone marrow, peripheral blood, spleens, and disease-inflicted tissues of different mouse models. GMP, granulocyte-monocyte progenitors. **b** Violin plots of the signature genes for neutrophil transcriptional states. **c** RNA velocity plot of neutrophil transcriptional states. **d** Pseudotime trajectory analysis of neutrophil transcriptional states. **e, f** UMAP plots of neutrophils in the diseased tissues **(e)** or the peripheral blood **(f)** of different mouse models. **g** Proportions of neutrophil transcriptional states in the bone marrow, peripheral blood, spleens, and diseased tissues of different mouse models. **h** Enrichment score heatmap of the gene sets of cytokine-related pathways in neutrophil transcriptional states. **i, j** C57BL/6 wild-type or *Ifngr1^-/-^* mice **(i)** or *Rag2^-/-^* and *Rag2^-/-^ Il2rg^-/-^* mice **(j)** were subjected to the ALI model. Total neutrophils in the peripheral blood, spleens, and lung tissues and their mean fluorescence intensity (MFI) of Cd274 expression were assessed by FACS. Mean ± SEM, * *p* < 0.05, ** *p* < 0.01, *** *p* < 0.001 (ANOVA test). **k** BALB/c wild-type or nude mice were utilized in the LCPNS-SIIN glioma model. Total neutrophils and Cd274^+^ neutrophils in the bone marrow, peripheral blood, spleens, and tumors were examined by FACS. Mean ± SD, * *p* < 0.05, ** *p* < 0.01 (ANOVA test). **l, m** C57BL/6 wild-type mice were subjected to ALI **(l)** or LCPNS-SIIN gliomas **(m)**. MFI of Cd244, Il4ra, or Cd101 expressed by neutrophils in the peripheral blood was quantified by FACS. Mean ± SD, ns not significant, ** *p* < 0.01 (Student’s *t-*test).

Of importance, in this comprehensive profiling of neutrophils under disease conditions, we observed two neutrophil clusters with unique immune characteristics. The first cluster included Neu_Acod1, Neu_Ccl4, and Neu_MHCII, which expressed the immunosuppressive signal *Cd274* (also known as Pd-l1; Fig. 1b) and myeloid recruitment-related chemokines *Ccl3* and *Ccl4* (Fig. S1a). Additionally, Neu_Acod1 and Neu_MHCII enriched the angiogenic factor *Vegfa*, and Neu_MHCII expressed the MHC class II genes such as *H2-Eb1* (Fig. 1b and Fig. S1a). In line with the previous studies showing the essential role of Cd274^+^ neutrophils in establishing the immunosuppressive tumor microenvironment of different cancers ^4,7,8^, we detected Neu_Ccl4 in all the examined peripheral or intracranial tumors (Fig. 1e, g). Also, consistent with the recent reports on the critical function of Vegfa^+^ or MHC-II^+^ neutrophils in antitumor immunity ^5,9,10^, Neu_MHCII was present in the peripheral or intracranial tumors except those in the immunodeficient NSG mice (Fig. 1e, g). Moreover, Neu_Ccl4 evidently occurred in the livers of HFD-fed mice and the lungs of ALI (Fig. 1e) while being almost absent in the peripheral blood (Fig. 1f), spleens (Fig. S2b), and bone marrow (Fig. S2c) of all the disease models. Supporting the validity of scRNA-seq results (Fig. 1e), FACS analyses confirmed the presence of both Neu_Ccl4 as Cd274^+^ MHCII^low^ and Neu_MHCII as Cd274^+^ MHCII^high^ neutrophils in the LLC lung tumors, while only Cd274^+^ MHCII^low^ neutrophils were in the lung tissues of ALI (Fig. S5a).

The Over Representation Analysis (ORA) of the gene sets of signaling pathways revealed that Neu_Ccl4 might be regulated by specific cytokines, e.g., tumor necrosis factor (TNF) or interferons (IFNs), as well as by hypoxia (Fig. 1h and Fig. S3). Meanwhile, Neu_MHCII was specifically related to the pathway of antigen presentation as expected (Fig. 1h and Fig. S3). In line with this notion, in the published scRNA-seq dataset (GSE202186) of mouse immune cells exposed to various cytokines ^11^, Neu_Ccl4 was also associated with the TNF and interferon signals (Fig. S4a). Indeed, we observed that the i*n vitro* treatment of neutrophils from mouse bone marrow with TNF-α, IFN-α, IFN-β, or IFN-γ were sufficient to elevate their expression of Cd274, as assessed by FACS (Fig. S4b). In parallel, RNA-seq analyses confirmed that those cytokines effectively induced the expression of the top 100 signature genes of scRNA-seq-defined Neu_Ccl4 (Fig. S4c). On the contrary, transforming growth factor beta (TGF-β) produced a minor effect, while interleukin 6 (IL-6) entirely failed to convert conventional neutrophils to the Neu_Ccl4 phenotype (Fig. S4b, c). We further examined the *in vivo* involvement of interferon signals in eliciting Neu_Ccl4. Cd274^+^ neutrophils could be robustly detected by FACS in the lung tissues of wild-type mice in the ALI model (Fig. S5c), but their appearance became diminished in the *Ifngr1^-/-^* mice (Fig. 1i). In addition, Cd274^+^ neutrophils was reduced in the lung tissues of *Rag2^-/-^* or *Rag2^-/-^ Il2rg^-/-^* mice subjected to ALI (Fig. 1j), implicating the possibility that lymphocyte-derived interferons would be involved in the Neu_Ccl4 induction. Similarly, FACS analyses showed Cd274^+^ neutrophils in the tumors but not the peripheral blood or spleens of wild-type mice in the LCPNS-SIIN glioma model (Fig. S5d), and their occurrence was significantly abolished in the nude mice that severely lacked T and B lymphocytes (Fig. 1k).

The second unique cluster of neutrophils, Neu_Cd244a, was characterized by the immunosuppressive signal *Cd244* ^12^ and type 2 immune response *Il4ra* (Fig. 1b and Fig. S1a). In contrast to the predominant presence of Neu_Ccl4 within diseased tissues, Neu_Cd244a appeared more enriched in the peripheral blood and spleens of specific disease conditions, e.g., ALI or LLC, but not other models, e.g., MC38 colorectal carcinoma, LCPNS glioma, or LCPNS-SIIN glioma (Fig. 1f, g and Fig. S2b). FACS analyses validated the presence of Neu_Cd244a in the peripheral blood of the ALI or LLC model, showing their increased expression of Cd244 and Il4ra but decreased maturation marker Cd101 (Fig. 1l and Fig. S5b). On the other hand, there was no significant change in Cd244, Il4ra, or Cd101 expressed by neutrophils in the peripheral blood of the LCPNS-SIIN glioma model (Fig. 1m), consistent with the absence of scRNA-seq-defined Neu_Cd244a in this disease scenario (Fig. 1f). Also, we compared Cd244^+^ neutrophils in the peripheral blood and Cd274^+^ neutrophils in the lung tissues of the ALI model by bulk RNA-seq analyses, verifying their distinct enrichment of *Cd244* and *Cd274* expression, respectively (Fig. S5e). To examine the potential immunosuppressive function of Neu_Cd244a, FACS-sorted Cd244^+^ neutrophils were co-cultured with CD8^+^ T cells. As a positive control of this *in vitro* assay, Cd274^+^ neutrophils triggered the up-regulation of Tim3 and Lag3 in Cd8^+^ T cells, the two key markers for their exhaustion ^13^, while Pd1 levels were mostly unaffected (Fig. S6a). Of importance, Cd244^+^ neutrophils could effectively elevate Cd8^+^ T cell expression of Tim3 and Lag3 (Fig. S6a). Moreover, RNA-seq analyses demonstrated that administration of an anti-Cd244 neutralizing antibody significantly increased the expression of a collection of pro-inflammatory cytokines and chemokines in the lung tissues of the ALI model (Fig. S6b).

In sum, this study has systematically profiled neutrophil transcriptional states and elucidated their common features in different mouse disease models, providing a valuable reference for future research on the pleiotropic functions of neutrophils in various pathophysiological contexts. Also, the identification of two distinct neutrophil clusters has implicated novel entry points for diagnostic strategies to stratify patients with specific disease conditions.

## Acknowledgments

This work was funded by the National Key Research and Development Program of China (#2022YFA1103900 to J.C.), the National Natural Science Foundation of China (#32125017 and #82441057 to J.Y.), the Ministry of Science and Technology of China (2021ZD0200800 to E.Y.), and the CAMS Innovation Fund for Medical Sciences (#2024-I2M-3-022 to J.C.). J.C. is also supported by the Chinese Institute for Brain Research and the Changping Laboratory.

## Author Contributions

D.W., Y.C., H.C., T.W., J.Z., and M.L. performed the experiments. D.W., M.D., W.W., H.Z., E.Y., J.Y., and J.C. analyzed the data. D.W., J.Y., and J.C. prepared the manuscript.

## Supplementary Materials and Methods

### Mouse information

All the experimental procedures were performed in compliance with the protocols approved by the Institutional Animal Care and Use Committee (IACUC) of the Chinese Institute for Brain Research or Peking University. Mice were maintained on the 12 h/12 h light/dark cycle (light period 7:00 am ∼ 7:00 pm) at ambient temperature (21°C ∼ 24°C) with water and standard chow diet available *ad libitum* unless otherwise specified.

C57BL/6 wild-type, BALB/c wild-type, and BALB/c nude mice were purchased from Charles River International. NSG (#005557) and *OT-1* (#003831) were from the Jackson Laboratory. *Ifngr1^-/-^* mice (#NM-KO-190432) on the C57BL/6 background were from Shanghai Model Organisms Center, and *Rag2^-/-^*mice (#C001324) or *Rag2^-/-^ Il2rg^-/-^* mice (#C001367) on the C57BL/6 background were from Cyagen Bioscience. 8- to 12- week-old female mice were utilized in the experiments unless otherwise specified.

### Mouse disease models

For the model of metabolic disorder, 5-week-old C57BL/6 male mice were fed with a high-fat diet containing 60% of calories from fat, 20% of calories from carbohydrates, and 20% of calories from proteins (Research Diets) for 24 weeks.

For the model of experimental autoimmune encephalomyelitis (EAE), myelin oligodendrocyte glycoprotein peptide (MOG_35-55_; Sangon Biotech) was dissolved in PBS and emulsified with an equal volume of complete Freund’s adjuvant (Chondrex) to achieve a final concentration of 1 mg/ml. This antigen emulsion was subcutaneously injected at four sites (50 μl per site) on the back of each mouse weekly for three weeks. Pertussis toxin (Sigma; 200 ng per mouse) was intraperitoneally administered on the first day and 48 hr after the first immunization. Tissues were harvested 45 days after the first MOG_35-55_ immunization.

For the model of acute lung injury (ALI), mice were briefly anesthetized with 3% isoflurane. 80 μg of lipopolysaccharide (LPS; Sigma) dissolved in 40 μl of sterile phosphate-buffered saline (PBS) was intranasally administered to each C57BL/6 female mouse. Tissues were harvested 48 hr after the LPS challenge. For the administration of the anti-Cd244 neutralizing antibody (clone OX-122, Biorbyt), each mouse was intraperitoneally injected with 20 μg antibody immediately after the LPS challenge.

For the model of Lewis lung carcinoma (LLC), 2 × 10^6^ LLC cancer cells suspended in 200 μl Leibovitz’s L-15 medium (Thermo Fisher Scientific) were inoculated into each mouse via intravenous injection. Tissues were harvested 30 days after the cell inoculation.

For the model of B16 melanoma or MC38 colorectal carcinoma, 1 × 10^6^ B16-OVA or MC38 cancer cells suspended in 100 μl Matrigel (Corning) were subcutaneously injected at the right flank of each mouse. Tissues were harvested 21 days after the cell inoculation.

For the model of intracranial glioma tumors, mice were anesthetized and head-fixed on the stereotactic frame. The head skin was shaved and prepared with iodine and 75% alcohol. A skin incision was made along the midline to expose the skull. The following coordinates for injection were measured relative to the bregma: anteroposterior = 0.5 mm, medial-lateral = 1.9 mm, and dorsal-ventral = −3.2 mm. A small hole was drilled through the skull at the target site, through which a Hamilton syringe needle (701N, Hamilton) was inserted into the brain. 5 × 10^4^ LCPNS or LCPNS-SIIN cancer cell lines were suspended in 5 μl Matrigel and delivered at the injection rate of 1.5 μl/min. The needle was kept in place for an additional 5 min to prevent the leakage of injected cells. Bone wax was used to cover the small hole in the skull to prevent the cells from overflowing, and the head skin was sutured. Tissues were harvested 21 days after the glioma inoculation.

### Cell lines and cultures

LCPNS or LCPNS-SIIN glioma cell lines were established as we recently reported ^1^. LCPNS or LCPNS-SIIN cells were cultured at 37°C under 5% CO_2_ / 5% O_2_ in Dulbecco’s Modified Eagle Medium/Nutrient Mixture F-12 (DMEM/F12 medium; Gibco) supplemented with 1% N2 (Gibco), 1% B-27 (Gibco), 1% GlutaMAX (Gibco), 100 U/ml penicillin, 100 μg/ml streptomycin, 16.6 mM D-glucose, 5 mM HEPES, 50 nM 2-mercaptoethanol (Sigma), 20 ng/ml EGF (Novoprotein), and 20 ng/ml FGF2 (Origene).

LLC, B16-F10, and MC38 cell lines were purchased from the Chinese National Infrastructure of Cell Line Resource and tested negative for mycoplasma. B16-OVA cancer cells were derived from B16-F10 cells by transducing with a lentivirus expressing full-length ovalbumin cDNA. LLC, B16-OVA, and MC38 cells were cultured 37°C under 5% CO_2_ in Dulbecco’s Modified Eagle Medium (DMEM; Gibco) supplemented with 10% fetal bovine serum (FBS; VISTECH), 100 U/mL penicillin, and 100 μg/mL streptomycin.

### Fluorescence-activated cell sorting

In light of published reports on the potential *ex vivo* activation of neutrophils ^2,3^, the FACS procedures were completed within 6 hr and on ice or at 4°C whenever necessary to minimize the impact on neutrophil transcriptional states.

The peripheral blood of each mouse was collected via retro-orbital bleeding into PBS containing 5 mM Na-EDTA (pH 8.0).

The spleen of each mouse was freshly dissected and directly mashed through a 70-μm cell strainer in Hank’s Balanced Salt Solution (HBSS) containing 3% heat-inactivated fetal bovine serum (HI-FBS; Sigma).

The tibias of each mouse were freshly dissected, and the bone marrow was flushed out with HBSS containing 3% HI-FBS and filtered through a 70-μm cell strainer. The skulls of each mouse were freshly dissected, and the dura mater partially connected was removed. The skull was cut into small pieces and incubated with 360° rotation in RPMI-1640 medium (Gibco) containing 3% HI-FBS and 10 mM HEPES at 37°C for 10 min to flesh out the skull bone marrow, and the cells were filtered through a 70-μm cell strainer.

The spinal cord of each mouse was freshly dissected and cut into small pieces on ice. The tissue was digested in an adequate volume of Accutase (BioLegend) at 37°C for 15 min and then mashed through a 70-μm cell strainer.

The lungs of each mouse were freshly dissected and cut into small pieces on ice.

The tissue was digested in RPMI-1640 medium containing 0.1 mg/ml Liberase TL (Roche), 20 μg/ml DNase I (Sigma), 10 mM HEPES, and 3% HI-FBS at 37°C for 15 min. The tissue was then mashed through a 70-μm cell strainer.

LLC, B16, or MC38 peripheral tumors were freshly dissected and cut into small pieces on ice. The tumor tissues were digested in RPMI-1640 medium containing 0.1 mg/ml Liberase TL, 20 μg/ml DNase I, 10 mM HEPES, and 3% HI-FBS at 37°C for 15 min. The tissues were then mashed through 70-μm cell strainers.

LCPNS or LCPNS-SIIN glioma tumors were freshly dissected and cut into small pieces on ice. The tumor tissues were digested in an adequate volume of Accutase at 37°C for 15 min and then mashed through 70-μm cell strainers.

Cell suspensions prepared from different tissues were centrifuged at 500 *g* for 5 min. The cells were re-suspended in ammonium-chloride-potassium (ACK buffer; Thermo Fisher Scientific) to lyse red blood cells. The cells were centrifuged again at 500 *g* for 5 min and re-suspended in HBSS containing 10 mM EDTA-Na (pH 8.0) and 2% HI-FBS for staining with the intended FACS antibodies and 7-AAD (Thermo Fisher Scientific). FACS antibodies utilized in this study included Cd45-PE (#103106, BioLegend), Cd45-PE-Cy7 (#103114, BioLegend), Cd11b-BV510 (#101263, BioLegend), Ly6G-BV421 (#127628, BioLegend), Ly6G-FITC (#127606, BioLegend), Ly6C-APC-Cy7 (#128026, BioLegend), Cd274-APC (#124312, BioLegend), MHCII-PE (#107607, BioLegend), Cd101-AF700 (#56-1011-82, eBioscience), Il4ra-AF488 (#53-1241-82, eBioscience), and Cd244a-PE-Cy7 (#133511, BioLegend).

The stained cells were processed on the BD LSRFortessa, and the FACS results were analyzed by FlowJo (https://www.flowjo.com). Alternatively, neutrophils (Cd45^+^ Cd11b^+^ Ly6G^+^ Ly6C^low^) were sorted on BD FACSAria for transcriptomic profiling.

### Single-cell RNA sequencing

Singleron Matrix Workflow: Single-cell suspensions (2 × 10^5^ cells/ml) in PBS were prepared and loaded onto a microwell chip using the Singleron Matrix Single Cell Processing System. Barcoding beads were collected from the chip, and mRNAs captured on beads underwent reverse transcription. The resulting cDNAs were amplified by PCR, fragmented, and ligated with sequencing adapters. Single-cell RNA-seq libraries were constructed following the manufacturer’s protocol for the GEXSCOPE Single Cell RNA Library Kit. The libraries were diluted to 4 nM and sequenced on the Illumina NovaSeq 6000 platform using 150 base pair (bp) paired-end reads.

SeekOne Workflow: Single-cell RNA-seq libraries were also generated using the SeekOne Digital Droplet Single Cell 3’ Library Preparation Kit (SeekGene). Single-cell suspensions were mixed with reverse transcription reagents and loaded into the sample wells of the SeekOne Chip S3. Barcoded hydrogel beads and partitioning oil were dispensed separately to generate droplets containing single cells. Reverse transcription was performed within the emulsion droplets, followed by droplet breaking to release cDNAs, which were subsequently purified and amplified. The amplified cDNAs were then processed through fragmentation, end-repair, A-tailing, and ligation to sequencing adapters. An indexed PCR was performed to amplify DNA corresponding to the 3’ polyA tails of genes, incorporating both the Cell Barcode and the Unique Molecular Index (UMI). The indexed libraries were purified using VAHTS DNA Clean Beads (Vazyme) and assessed for quality and concentration using the Qubit (Thermo Fisher Scientific) and the Bio-Fragment Analyzer (BiOptic). The libraries were sequenced on the Illumina NovaSeq X Plus platform with 150 bp paired-end reads.

### Single-cell RNA sequencing data processing

Data preprocessing was performed according to the manufacturer’s reference instructions. The reference mouse genome version used for sequence alignment was mm10. For quality control, we filtered out genes expressed in <10 cells, as well as cells expressing gene numbers <200, mitochondrial reads >10%, and ribosome genes >20%. We used the Solo in scvi-tools to remove doublet using the top 2,000 variant genes and default parameters.

To integrate scRNA-seq data from multiple samples, we adopted the methods of recent scRNA-seq studies of neutrophils ^1,4–6^. We utilized single-cell Variational Inference (scVI) (https://docs.scvi-tools.org/en/stable/tutorials/) from the scvi-tools package. Data clustering was performed using Scanpy (https://github.com/scverse/scanpy), and the resulting H5ad files were converted to Seurat objects using Sceasy (https://github.com/cellgeni/sceasy). The analysis was conducted in RStudio with R v4.2.2 (https://cran.r-project.org/) and Seurat v4 (https://satijalab.org/seurat/articles/get_started.html). Marker genes for each cluster were identified and visualized using the FindAllMarkers function in Seurat. Cell type annotation was initially performed using a combination of ScType (https://sctype.app) and PanglaoDB (https://panglaodb.se/index.html). For detailed characterization of neutrophil subtypes, we referenced previously published studies ^1,7,8^. To ensure balanced visualization across samples, during the UMAP embedding process stratified by tissue types or disease conditions, we down-sampled each sample to a maximum of 5,000 cells.

### Integrated analysis of RNA velocity, pseudotime, and gene regulation

RNA velocity analysis was performed using scVelo (https://scvelo.readthedocs.io/en/stable/) with default parameters. The input data for scVelo was generated using the Python implementation of Velocyto. Pseudotime trajectory analysis was conducted with Monocle 3 (https://cole-trapnell-lab.github.io/monocle3/) utilizing its default settings.

To investigate gene regulatory networks, we employed pySCENIC (v0.12.0) in a Docker environment (https://hub.docker.com/r/aertslab/pyscenic). Transcription factors differentially expressed across cell types were identified based on AUC scores and visualized using ComplexHeatmap (https://www.bioconductor.org/packages/devel/bioc/html/ComplexHeatmap.html) and ClusterProfiler (https://bioconductor.org/packages/release/bioc/html/clusterProfiler.html).

Gene set enrichment analysis was conducted using the Python version of DecoupleR (v1.7.1) (https://github.com/saezlab/decoupler-py) in combination with Scanpy (v1.10.2). The Over Representation Analysis (ORA) method was applied to gene sets from MSigDB (v7.4) (https://www.gsea-msigdb.org/gsea/msigdb), with a focus on identifying cytokine-associated gene sets.

### Mapping cytokine datasets to neutrophil subtypes

Raw FASTQ files containing single-cell transcriptomic profiles of immune cells exposed to various cytokines were obtained from the Gene Expression Omnibus (GEO) database under accession number GSE202186 ^9^. Sequencing reads were aligned to the mouse reference genome mm10, and transcriptomic count matrices were generated using the CellRanger pipeline (v7.2). For the hashtag library, FASTQ files were processed with CITE-seq-Count (v1.4.3) (github.com/Hoohm/CITE-seq-Count). Gene expression data were matched with hashtag information using the HTODemux function in the Seurat R package (v5.1.0) (https://satijalab.org/seurat/). Cells classified as multiplets (e.g., those with multiple hashtags) were excluded from downstream analysis.

Quality control was performed using the Scanpy package (v1.10.3). Cells were retained if they expressed <6,000 genes and mitochondrial gene contents <20%. Gene expression matrices were normalized by scaling each cell’s gene expression to its total expression, multiplying by a scaling factor of 10,000, and applying a log transformation. For dimensionality reduction, the top 2,000 most variable genes were selected. Principal component analysis (PCA) was then used to denoise the data and reduce it to a lower-dimensional representation, retaining the top 40 principal components that were used for global clustering and visualization with UMAP.

Neutrophils were identified at the cluster level based on UMAP clustering results and the expression of canonical marker genes, including *S100a8*, *Cxcr2*, *Itgam*, *Mmp9*, and *Csf3r*. Cytokine information for each neutrophil was inferred from the corresponding hashtags. Finally, the Seurat R package was used to map neutrophil clusters to neutrophil subtypes defined in the current study, enabling the identification of cytokines potentially regulating each subtype.

### Bulk RNA sequencing

Total mRNAs of neutrophils were extracted and reverse-transcribed using a template-switching oligo (TSO; BGI Genomics). cDNAs then underwent pre-amplification and tagmentation to add sequencing adapters, and the index PCR was performed to incorporate sample-specific barcodes. The libraries were purified using VAHTS DNA Clean Beads (Vazyme) and assessed for quality and concentration using the Qubit (Thermo Fisher Scientific) and the Bio-Fragment Analyzer (BiOptic). The libraries were sequenced on the MGI DNBSEQ-T7 platform using 150 bp paired-end reads. Sequencing files were aligned to the reference mouse genome using STAR (v2.7.11a) (https://code.google.com/archive/p/rna-star/). Gene expression levels were quantified with FeatureCounts included in the Subread package (https://subread.sourceforge.net/). Differential gene expression analysis was performed using edgeR (https://bioconductor.org/packages/release/bioc/html/edgeR.html). Pathway enrichment analysis was conducted using the fgsea package (https://bioconductor.org/packages/release/bioc/html/fgsea.html), with results visualized through ggplot2 (https://ggplot2.tidyverse.org) for clear and effective representation of enriched pathways from MSigDB (v7.4).

### *In vitro* treatments

Neutrophils were FACS-sorted from the tibial bone marrow of C57BL/6 wild-type mice and cultured in RPMI-1640 medium supplemented with 10% FBS, 100 U/mL penicillin, and 100 μg/mL streptomycin in 6-well plates (1 × 10^6^ neutrophils per well). Neutrophils were *in vitro* treated with a final concentration of 20 ng/ml IL-6 (PeproTech), TGF-β (Novoprotein), TNF-α (Novoprotein), IFN-α (Sinobiological), IFN-β (Sinobiological), or IFN-γ (Novoprotein) for 18 hr. *In vitro* treated neutrophils were then examined by FACS or bulk RNA-seq analyses.

The co-cultures of neutrophils and Cd8^+^ T cells were performed as we recently reported ^1^. The spleen and lymph nodes of OT-1 mice were freshly dissected and mashed through 70-μm cell strainers in HBSS containing 3% HI-FBS. The cells were centrifuged at 500 *g* for 5 min and re-suspended in ACK buffer to lyse red blood cells. The cells were centrifuged again at 500 *g* for 5 min and re-suspended in RPMI 1640 medium supplemented with 10% HI-FBS, 2 mM glutamine, 55 μM 2-mercaptoethanol, 1 mM sodium pyruvate, 100 U/ml penicillin, and 100 μg/ml streptomycin. After loading with 1 mM OVA_257-264_ peptide (SIINFEKL) at 37°C for 1 hr, the cells were centrifuged at 500 *g* for 5 min, re-suspended in culture medium, and cultured at 37°C overnight. The activated OT-1 Cd8^+^ T cells (Cd45^+^ Cd3^+^ Cd4^-^ Cd8^+^ NK1.1^-^) were then FACS sorted. In parallel, conventional neutrophils from the tibial bone marrow of C57BL/6 wild-type mice and Cd244^+^ neutrophils from the peripheral blood or Cd274^+^ neutrophils from the lung tissues of C57BL/6 wild-type mice in the ALI model were FACS sorted. 5 × 10^4^ OT-1 Cd8^+^ T cells were co-cultured with 5 × 10^4^ neutrophils for 6 hr before their expression of specific exhaustion markers were analyzed by FACS.

### Data availability

The scRNA-seq and RNA-seq data obtained in this study have been deposited at the National Genomics Data Center (https://ngdc.cncb.ac.cn/) under the accession number CRA020963.

### Statistical analysis

Statistical analyses were performed by GraphPad Prism 9.5.0 (http://www.graphpad.com/scientific-software/prism). All the sample points (n) shown in the figures represent biological replicates (i.e., mice or cell preparations). The statistical test description and *p* values can be found in the figure legends where appropriate.

## Supplementary Table

**Table S1.**
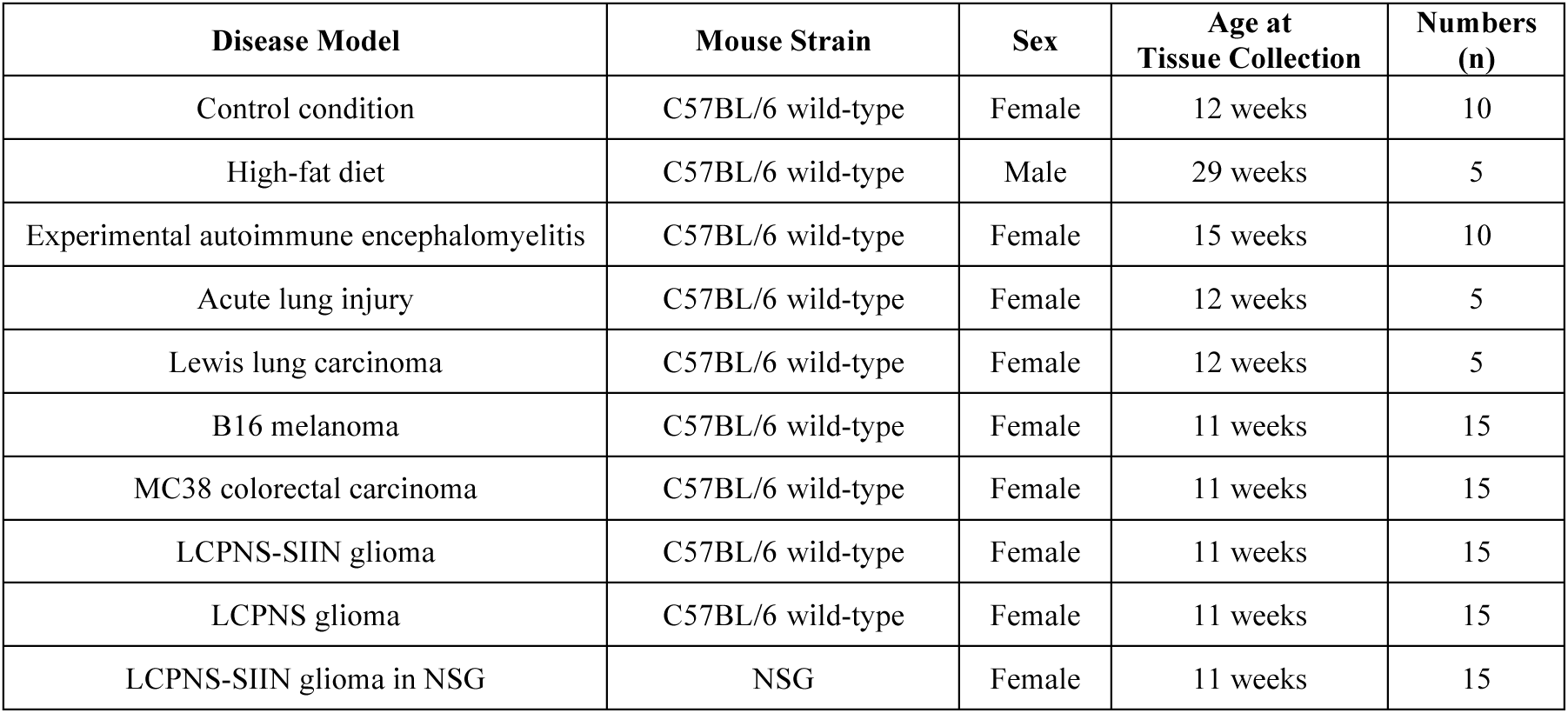
Mouse Information for scRNA-seq Analyses of Neutrophils.

## Supplementary Figures

**Figure S1.**
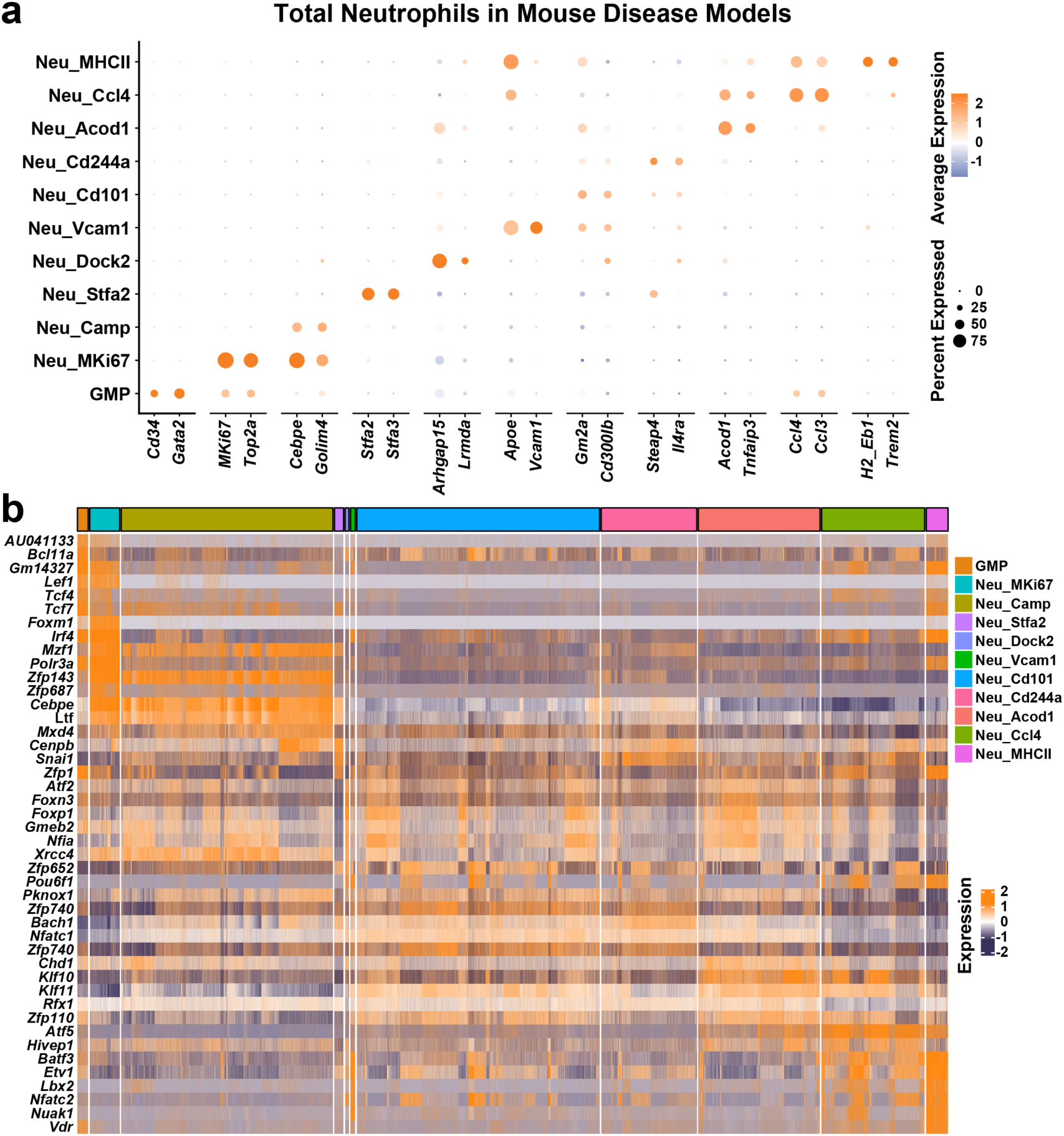
Signature genes of neutrophil transcriptional states. **a** Dot plot of two top marker genes for each neutrophil transcriptional state defined in the pooled scRNA-seq dataset of mouse disease models **b** Heatmap of SCENIC binary regulon activities in neutrophil transcriptional states.

**Figure S2.**
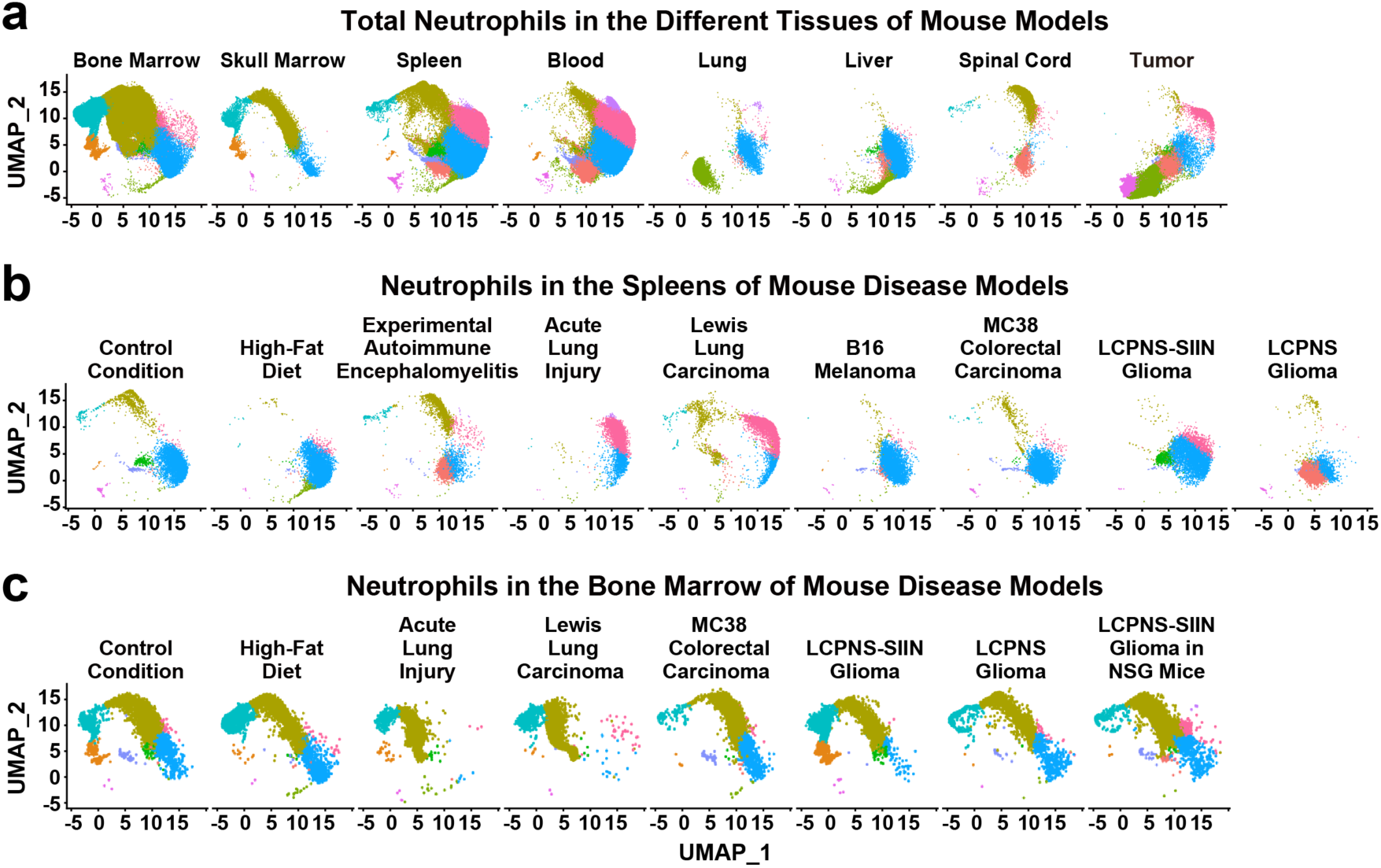
Neutrophil transcriptional states in different tissues and disease conditions. **a** UMAP plots of neutrophil transcriptional states in different tissues defined in the pooled scRNA-seq dataset of mouse disease models. **b, c** UMAP plots of neutrophil transcriptional states in the spleens **(b)** or bone marrow **(c)** of different disease models.

**Figure S3.**
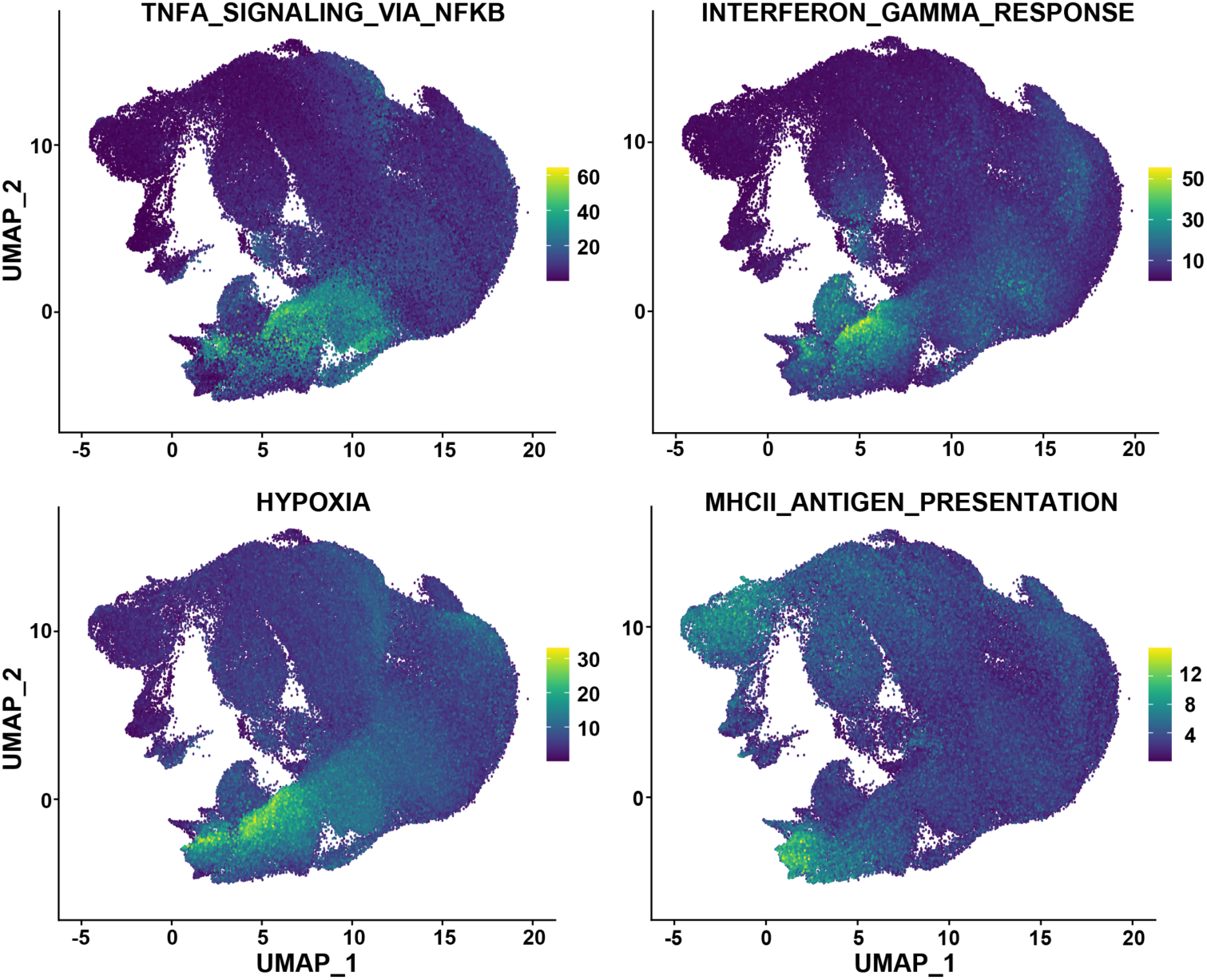
Enrichment of signaling pathways in neutrophil transcriptional states. Enrichment scores for the gene sets of specific signaling pathways calculated by the Over Representation Analysis are projected onto the UMAP plots of neutrophil transcriptional states defined in the pooled scRNA-seq dataset of mouse disease models.

**Figure S4.**
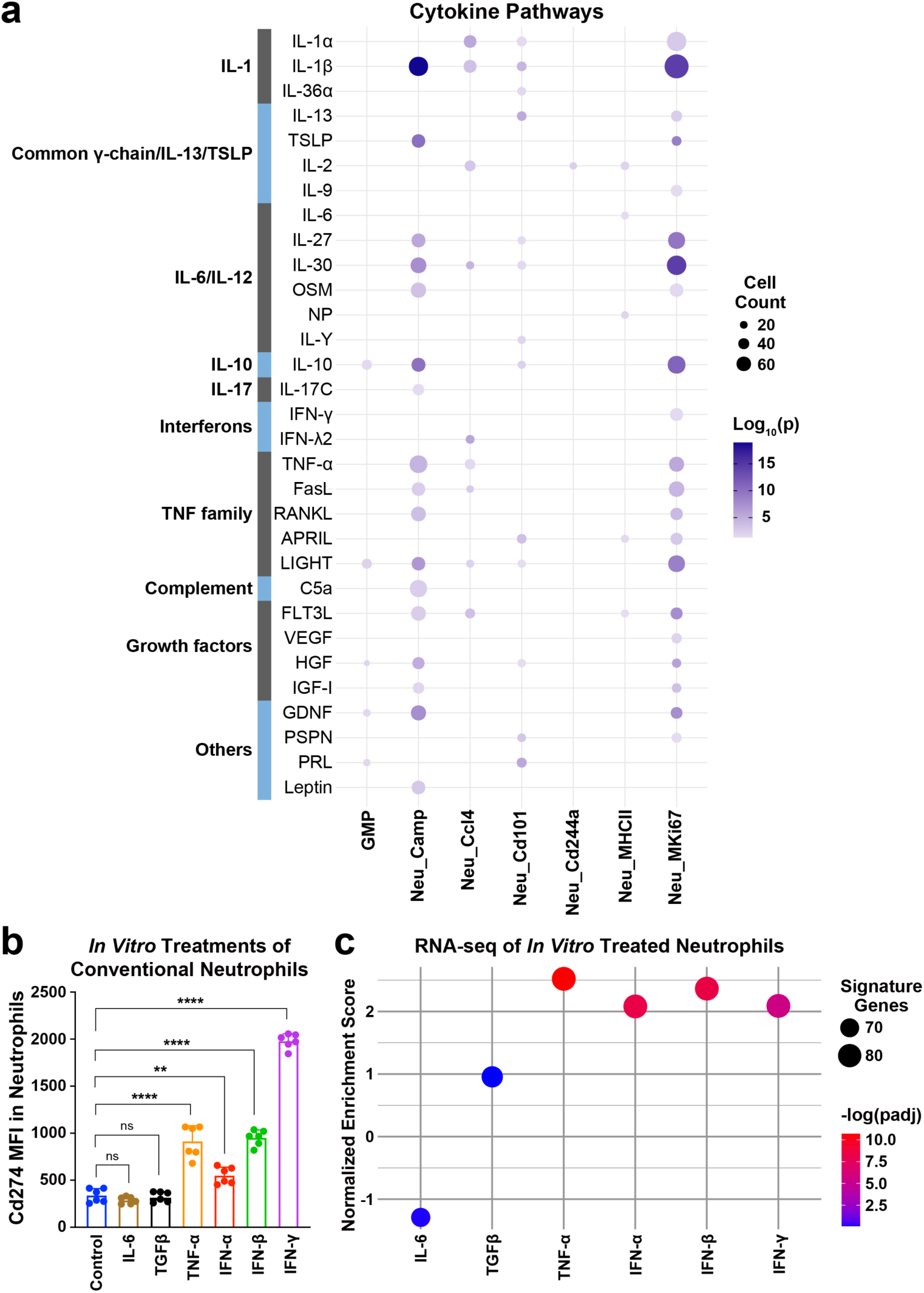
Cytokine signals for the induction of neutrophil transcriptional states. **a** Published scRNA-seq dataset of immune cells exposed to various cytokines (GSE202186) was re-analyzed. Neutrophils in the published dataset were mapped to specific transcriptional states defined in the current study. **b, c** Neutrophils were FACS-sorted from the tibial bone marrow of C57BL/6 wild-type mice and *in vitro* treated with the indicated cytokines. **b** Mean fluorescence intensity (MFI) of Cd274 expression in neutrophils was determined by FACS. Mean ± SD, ns not significant, ** *p* < 0.01, **** *p* < 0.0001 (ANOVA test). **b** *In vitro* treated neutrophils were profiled by bulk RNA-seq. Normalized enrichment scores of the top 100 marker genes of Neu_Ccl4 are shown.

**Figure S5.**
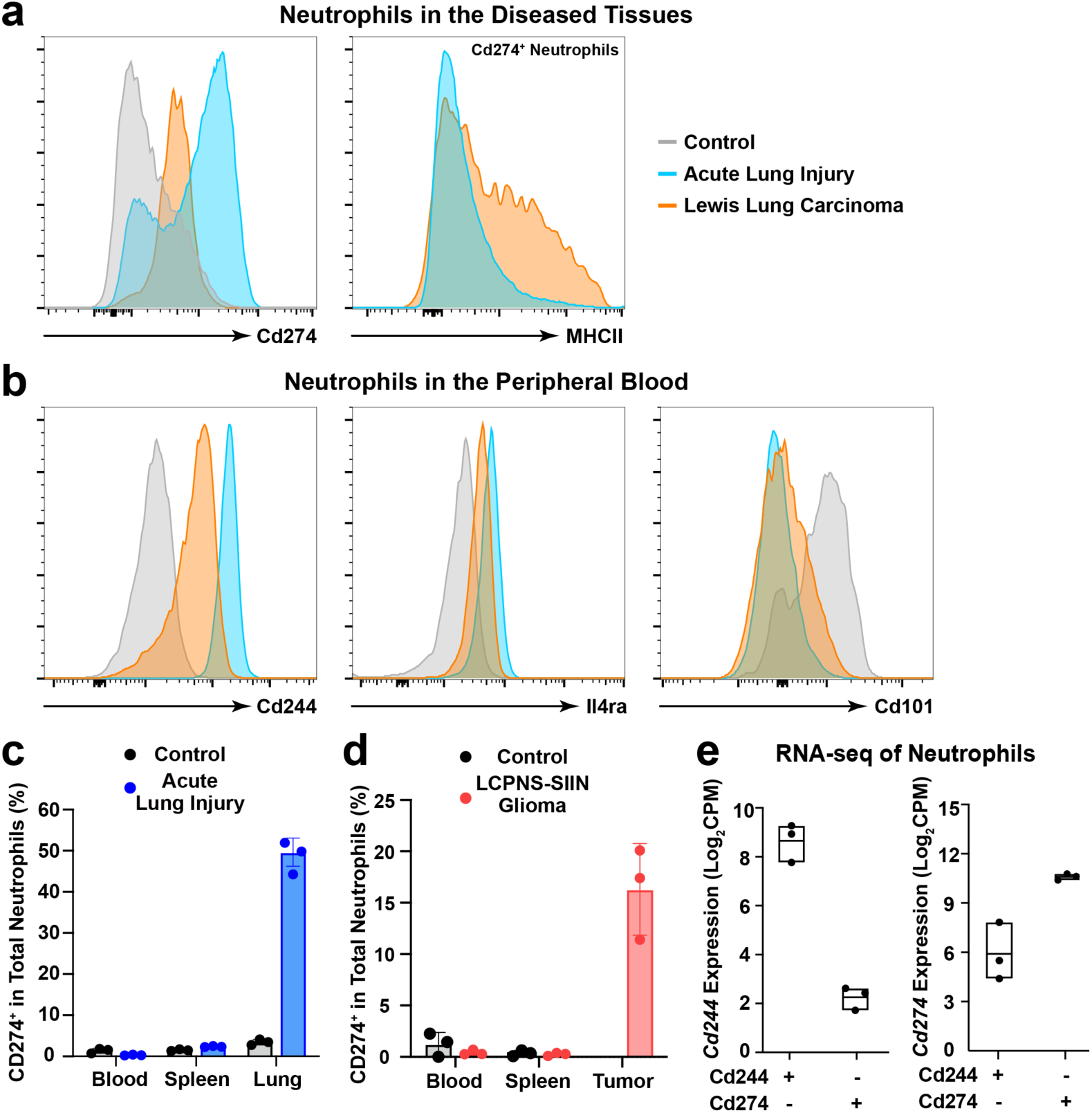
Characterization of neutrophil transcriptional states. C57BL/6 wild-type mice were subjected to the indicated disease models. **a** Cd274 and MHCII expression by total neutrophils in the lung tissues of control mice or the ALI model or the lung tumors of the LLC model were examined by FACS. Representative histograms are shown. **b** Cd244, Il4ra, and Cd101 expression by total neutrophils in the peripheral blood of control mice, the ALI model, or the LLC model were assessed by FACS. Representative histograms are shown. **c** Cd274^+^ neutrophils in the peripheral blood, spleens, and lung tissues of control mice or the ALI model were quantified by FACS. Mean ± SD. **d** Cd274^+^ neutrophils in the peripheral blood, spleens, and intracranial tumors of control mice or the LCPNS-SIIN glioma model were determined by FACS. Mean ± SD. **e** Cd244^+^ neutrophils from the peripheral blood and Cd274^+^ neutrophils from the lung tissues of the ALI model were profiled by bulk RNA-seq. Expression levels of *Cd244* and *Cd274* are presented as box plots.

**Figure S6.**
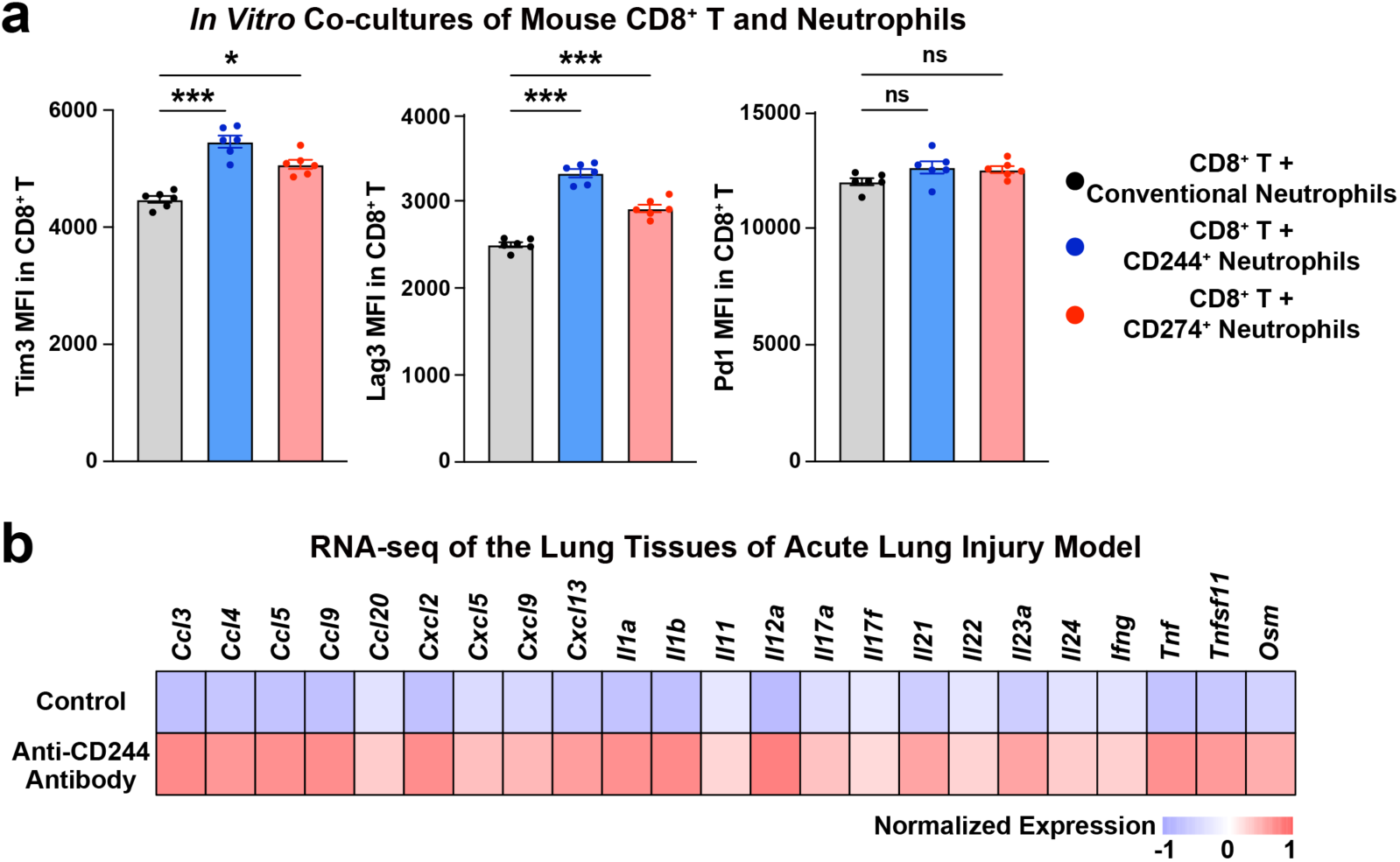
Immunosuppressive function of Cd244^+^ neutrophils. **a** Cd244^+^ neutrophils from the peripheral blood or Cd274^+^ neutrophils from the lung tissues of C57BL/6 wild-type mice in the ALI model were *in vitro* co-cultured with OT-1 Cd8^+^ T cells. The co-culture with conventional neutrophils from the tibial bone marrow of C57BL/6 wild-type mice was included as the control condition. Mean fluorescence intensity (MFI) of Tim3, Lag3, or Pd1 expression by Cd8^+^ T cells was examined by FACS. Mean ± SEM, ns not significant, * *p* < 0.05, *** *p* < 0.001 (ANOVA test). **b** C57BL/6 wild-type mice were treated with an anti-Cd244 neutralizing antibody immediately after the ALI model. The lung tissues were profiled by bulk RNA-seq (five mice per condition). Average expression of pro-inflammatory cytokines and chemokines up-regulated in the anti-Cd244 condition are shown.

